# The dynamics of protein localisation to restricted zones within *Drosophila* mechanosensory cilia

**DOI:** 10.1101/2022.03.02.482694

**Authors:** Wangchu Xiang, Petra zur Lage, Fay G. Newton, Guiyun Qiu, Andrew P. Jarman

**Author notes:** These authors contributed equally to this work (Wangchu Xiang, Petra zur Lage, Fay G. Newton).

## Abstract

The *Drosophila* chordotonal neuron cilium is the site of mechanosensory transduction. The cilium has a 9+0 axoneme structure and is highly sub-compartmentalised, with proximal and distal zones harbouring different TRP channels and the proximal zone axoneme also being decorated with axonemal dynein motor complexes. The activity of the dynein complexes is essential for mechanotransduction. We investigate the localisation of TRP channels and dynein motor complexes during ciliogenesis. Differences in timing of TRP channel localisation correlate with order of construction of the two ciliary zones. Dynein motor complexes are initially not confined to their target proximal zone, but complexes beyond the proximal zone are later cleared, perhaps by retrograde transport. Differences in transient distal localisation of ODAs and IDAs are consistent with previous suggestions from unicellular eukaryotes of differences in processivity during intraflagellar transport. Stable localisation depends on the targeting of their docking proteins in the proximal zone. For outer dynein arm (ODA) complexes, we characterise an ODA docking complex (ODA-DC) that is targeted directly to the proximal zone. Interestingly, the subunit composition of the ODA-DC in chordotonal neuron cilia appears to be different from the predicted ODA-DC in *Drosophila* sperm.

## INTRODUCTION

Cilia are compartmentalised organelles, and their growth (ciliogenesis) requires mechanisms for transport of cilium-resident proteins and protein complexes into the cilium and their incorporation into the ciliary membrane or onto the microtubular axoneme. Ciliogenesis proceeds by growth at the tip of the cilium. In general, at the basal body/transition zone the majority of ciliary proteins are loaded as cargoes onto ‘trains’ for transport along the axonemal microtubules by a dedicated process called intraflagellar transport (IFT) (Kozminski et al., 1993, 1995; Rosenbaum and Witman, 2002). Anterograde movement of IFT trains from the base to the tip of the flagellum/cilium is powered by kinesin-2 motors interacting with IFT-B proteins; and retrograde movement from the tip to the base by IFT dynein-2 motors with IFT-A proteins. Most cargoes appear to be transported to the growing ciliary tip before being released for incorporation into the ciliary structure (Johnson and Rosenbaum, 1992), although some cargoes may be released during anterograde transport itself.

In motile cilia, major cargoes of IFT are the specialised axonemal dynein motor complexes that power ciliary movement. These form Inner and Outer Dynein Arms (IDAs and ODAs) (Fig. 1A). The multicomponent motor complexes are very large (1–2 MDa in size), with subunits including the force-generating heavy chains (HC), scaffolding intermediate and light-intermediate chains (IC, LIC), as well as light chains (LC, < 30 kDa) that regulate protein interaction and microtubule-anchoring (King, 2016) (Fig. 1A). In our current understanding, especially from studies in unicellular organisms such as the biflagellate *Chlamydomonas reinhardtii,* dynein motor complexes are pre-assembled within the cytoplasm before transport into the cilium and docking on the axoneme (Fowkes and Mitchell, 1998). Pre-assembly requires dedicated co-chaperone assembly factors (generally known as DNAAFs) (Desai et al., 2018; Horani and Ferkol, 2013) (Fig. 1B). These pre-assembled complexes must move past the transition zone barrier at the base of the cilium and are then transported through the cilium by IFT trains, to which they bind with the help of adaptor proteins (Dai et al., 2018; Viswanadha et al., 2014) (Fig. 1B). Subsequent events are less clear and may vary between complexes. IFT trains may take motor cargoes to the tip of flagellum, where cargoes undergo ‘maturation’ and release (Viswanadha et al., 2014), or they may be released during anterograde transport (Dai et al., 2018; Wren et al., 2013). The complexes are then thought to diffuse locally onto their microtubule anchoring sites, where their periodic binding is guided and stabilised by pre-bound ‘docking proteins’. For ODAs, this consists of a three-subunit ODA docking complex (ODA-DC) (Wakabayashi et al., 2001); for IDAs and other motility complexes, several proteins participate in docking, including a ‘96 nm molecular ruler’ formed of two coiled-coil proteins (Ma et al., 2019) (Fig. 1A).

**Figure 1.**
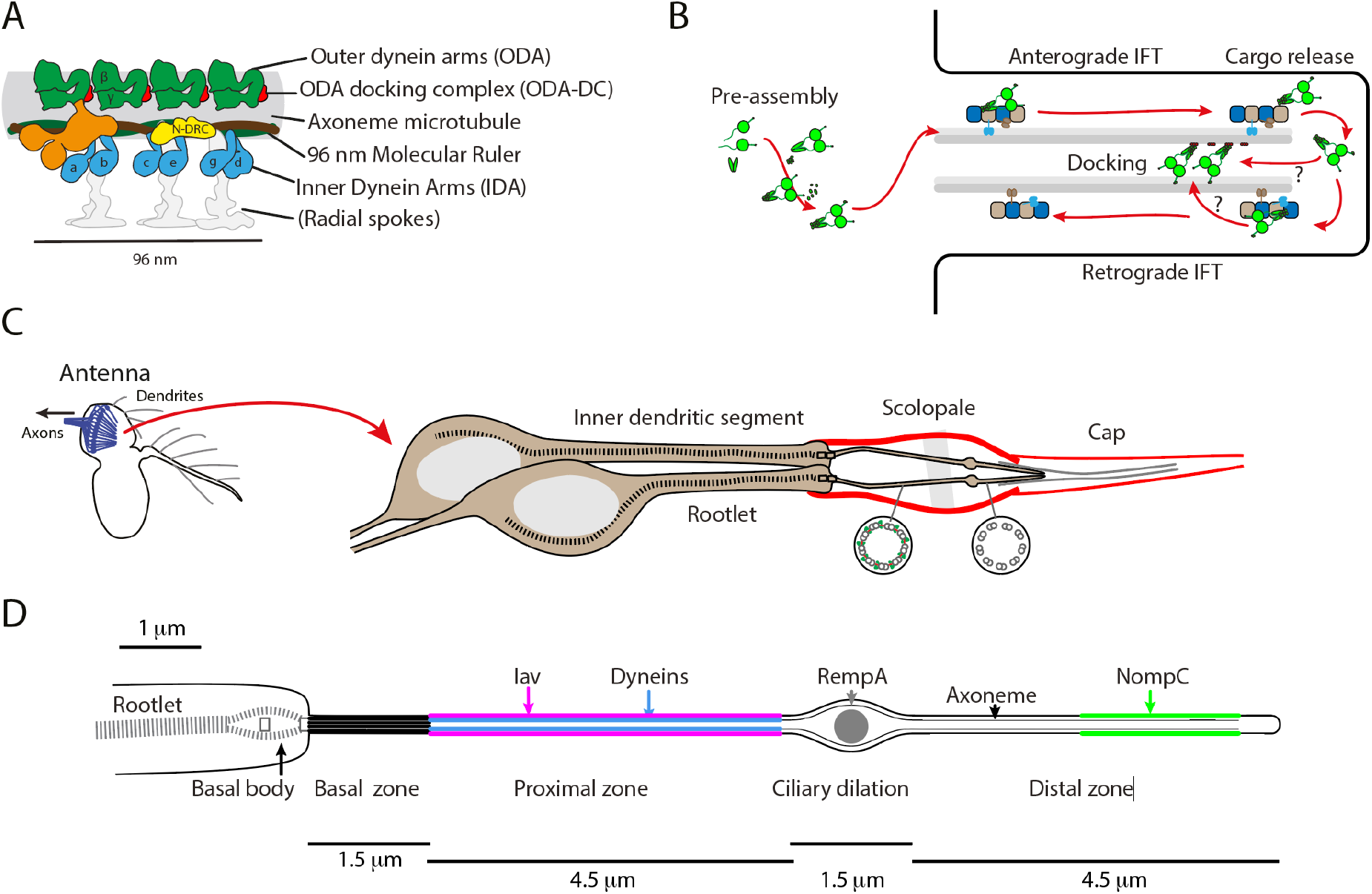
Axonemal dyneins and the chordotonal neuron cilium. (A) Schematic of the axonemal protein complexes of motile cilia. The repeating unit of a single microtubule doublet is represented. Chordotonal cilia have a 9+0 axoneme and so lack the radial spokes (greyed out). N-DRC: nexin-dynein regulatory complex. Based on (zur Lage et al., 2019). (B) Schematic outline of generalised axonemal dynein assembly and the intraflagellar transport (IFT) pathway. (C) The structure of chordotonal scolopidia of Johnston’s organ (JO) in the *Drosophila* antenna. JO is housed in the second antennal segment. Each ‘unit scolopodium’ comprises 2-3 mechanosensory chordotonal neurons. Each dendrite terminates in single 9+0 cilium. The cilia are bathed in lymph within a scolopale structure and are attached at their tips to a cap structure, enabling the transmission of mechanical force to the cilia. (D) Schematic of the adult chordotonal neuron cilium (outer dendritic segment). The cilium is divided into a ‘motile’ proximal zone and a ‘sensory’ distal zone, between which is the ciliary dilation (CD). Axonemal dyneins and TRPV channels (Iav/Nan) are confined to the proximal zone wherease TRPN channels (NompC) are in a region of the distal zone. Several proteins reside at the ciliary dilation, including the IFT-A protein, RempA. Lengths are approximate, based on electron micrographs of adult JO cilia.

The dynein assembly pathway is thought to be conserved in humans and other metazoans, including *Drosophila* (zur Lage et al., 2019). The pathway is medically important because defects in it are a frequent cause of the inherited disease, primary ciliary dyskinesia (PCD), characterised by reduced or absent ciliary motility (Mitchison and Valente, 2017). However, while such studies have identified conserved components, there are also differences in metazoans. For instance, the ODA-DC seems to have a different composition from *Chlamydomonas* (Knowles et al., 2013; Onoufriadis et al., 2014). Moreover, mechanistic evidence in metazoans for dynein motor transport and docking is sparse.

It is increasingly apparent that ciliary proteins can be located to different regions of a cilium/flagellum for functional reasons. The mechanisms by which such sub-compartmentation is achieved during assembly are poorly known. A striking example is found in *Drosophila* in the ciliated dendrite of auditory/proprioceptive sensory neurons (chordotonal neurons) of the antennal sensory organ known as Johnston’s Organ (Fig. 1C). These specialised 9+0 cilia are notable for two reasons: (1) They have the features of motile cilia, including axonemal dynein motor complexes (zur Lage et al., 2019) the function of which is critical to their mechanotransduction mechanism (Karak et al., 2015; zur Lage et al., 2021); (2) They have structurally distinct zones with specialised functions: a sensory ‘distal zone’ closely connected with an external ciliary cap and containing the mechanosensory TRP channel, NompC, and a motile ‘proximal zone’ containing heteromeric TRPV channels (Inactive/Nanchung or lav/Nan) as well as the axonemal dynein motors (Fig. 1D). These zones are separated by a ‘ciliary dilation’ of unclear function. Chordotonal neuron cilia thus serve as a model to address two questions: how are dynein motors assembled onto axonemes? How are these and other proteins targeted to zones within cilia?

For the latter question, localisation of the TRP channels has been extensively studied. Based on mutant analyses of IFT-A proteins *(rempA* and *Oseg4)* and IFT dynein *(beethoven),* retrograde IFT plays a role in TRP channel entry into the cilium and correct localisation to distinct zones (Eberl et al., 2000; Lee et al., 2008). Since the ciliary dilation is also defective in these mutants, and IFT-A protein RempA accumulates at the ciliary dilation that separates the zones (Fig. 1D), it is suggested that the ciliary dilation is required for correct TRP targeting. But an alternative interpretation is that these retrograde transport genes regulate TRP targeting and ciliary dilation structure in parallel. As with other ciliary membrane proteins, Iav and NompC localisation require a Tulp protein (dTulp/king tubby) as an adaptor linking these cargoes to IFT (Mukhopadhyay and Jackson, 2011; Park et al., 2013). As in other organisms, release of TRP cargo from dTulp appears to be regulated by PIP levels in the cilium (Park et al., 2013, 2015). In contrast to TRP channels, nothing is known about how dynein motors are transported and then released and localised specifically to the proximal zone of chordotonal cilia.

To investigate protein localisation in chordotonal cilia, we first characterise TRP channel localisation during the time course of chordotonal ciliogenesis. Using this as a guide, we then demonstrate qualitative differences in how ODA and IDA motors become localised over time, which reflect prior differences in localisation of ODA-DC and the molecular ruler complex, respectively. We show that ODA is targeted directly to the proximal zone via an ODA-DC containing conserved Ccdc114 and Ccdc151 subunits. In contrast, IDAs appear to be transported to the ciliary tip before stable docking in the proximal zone. This mode is reflected by IDA docking protein, Ccdc39. Our data are consistent with a model in which ‘outer’ axonemal proteins (ODA and ODA-DC) are released from IFT trains during transport for local axonemal binding, while ‘inner’ axonemal proteins (including IDAs and their docking factors) are released at the ciliary tip to gain access inside the growing axoneme. In addition, we discuss the finding that ODA-DC appears to be an entirely different structure in chordotonal cilia compared with sperm flagella.

## METHODS

### Fly stocks

All fly strains were reared on a standard cornmeal agar medium at 25°. The following stocks were obtained from the Bloomington *Drosophila* Stock Center (Indiana University, Bloomington, IN): y[1]w[*]P{y[t7.7]=nos-phiC31\int.NLS}X; P{y[t7.7]=CaryP}attP40 (#79604), *y[1]M{RFP3xP3.PB]GFP[E.3xP3]=vas-Cas9}ZH-2A w[1118]/FM7c* (#51323), *w1118* (#3605) and P{w[=mC]=UAS-Dcr-2.D}1, w[1118]; Pin[1]/CyO P (#24644). CG14127-GFP, CG4329-GFP, CG6652-GFP stocks were gifts from Bénédicte Durand. The *sca-Gal4* line was a gift from M. Mlodzik/N. Baker (Baker et al., 1996). The RNAi lines were obtained from the Vienna *Drosophila* Resource Centre (VDRC) (Dietzl et al., 2007): UAS-*CG14127* RNAi (106258), UAS-*CG3723* RNAi (108658), UAS-dTulp RNAi (29111) and control (KK) line (60100). For the dTulp RNAi crosses, the pupae were raised at 28°C.

### *mVenus* and *mCherry* fusion gene constructs

*mVenus* fusion constructs for *CG6971/IDA-Dnali1, CG8800/ODA-Dnal1, CG17387/Ccdc39* and *CG14905/Ccdc114* were designed to fuse the coding region of mVenus in frame at the 3’ end of the coding region of the target gene (primers in Table S1). The target gene fragment was amplified to include the entire ORF and sufficient upstream regulatory region to ensure the spatial-temporal pattern of expression from its own promoter (as judged by presence of conserved binding sites for transcription factors Rfx and Fd3F, (Newton et al., 2012)). An *att*B-flanked gene fragment was amplified by PCR from genomic DNA. Using the Gateway® two-step recombination system (Life Technologies), the fragment was first inserted into pDONR221 in the BP reaction and then incorporated into a modified pBID-UASC-GV vector (Wang *et al.,* 2012) using the LR reaction mix. The UAS site deletion was generated performing single primer mutagenesis using Pfu Turbo Cx DNA polymerase (Agilent). The pBID-GmCherry vector was constructed by replacing the mVenus of the pBID-GV vector by the mCherry sequence. This was achieved by digesting the pBID-GV vector with *XhoI* and *XbaI,* followed by Gibson cloning (NEBuilder HiFi). The *CG6971/IDA-Dnali1* (Moore et al., 2013) and the above mentioned *CG8800/ODA-Dnal1* pDONR221 BP plasmids were repurposed for the cloning into the LR destination plasmid pBID-GmCherry. Transformant fly lines were generated by microinjection into syncytial blastoderm embryos of the attP40 landing site line using nos-PhiC31 integrase.

### Engineering of *CG14905* Crispr/Cas9 deletion mutant

A *CG14905 CRISPR/Cas9* mutant was constructed by a mini-white gene substitution according to zur Lage et al. (zur Lage et al., 2018). The guide RNA plasmid and the homology arm plasmid were subsequently injected together into the Cas9 injection line vasa::Cas9 to generate the transformant line (primers in Table S1).

### *Drosophila* embryonic microinjection

Purified plasmids (GeneJET Plasmid Miniprep Kit, Thermo Scientific K0503) were microinjected into embryos of *nos/attP40* flies for mVenus fusion gene construct injection and *nos-Cas9* flies for CRISPR mutagenesis, which contains the 1:1 mix solution of the gRNA expression construct and the homology arm expression construct. Single fly PCRs were used to confirm correct substitution of CG14905 with mini-*white*. Two pairs of primers were used, each containing one primer inside *miniwhite* gene sequence and another primer inside upstream/downstream homology arm sequence.

### Genomic DNA extraction from adult fruit flies

50 flies from the same line were frozen in lysis buffer (100mM Tris-HCl [pH9], 100mM EDTA, 1% SDS) at −20°C. After thawing, the sample was homogenized and incubated the sample at 70°C for 30min. Potassium acetate (8M) was then added and the sample was incubated on ice. After centrifugation, the supernatant was transferred, and nucleic acids precipitated using isopropanol. After redissolving the pellet, the solution was extracted with Phenol-chloroform (Sigma #P3803) and precipitated with ethanol. DNA was redissolved in TE.

### Immunohistochemistry for pupal antennae

White puparia were aged for the appropriate period of time at 25°C. The anterior puparium case was opened up and removed to uncover the pupal head region and the whole pupa was fixed in 3.7% formaldehyde for 30 minutes, followed by washing in PBT (0.3% Triton X-100). Subsequently, the antennae were dissected off, collected in a 1.5 ml tube and blocked overnight in 3% BSA (Roche) before incubating in primary antibody (3% BSA in PBT) at 4°C for 2 days. After washing in PBT, antennae were incubated in secondary antibodies at 4°C overnight. They were mounted in a 2.5% n-Propyl gallate/glycerol mix. The following primary antibodies were used: Goat anti-GFP (1:500: Abcam, ab6673), Rabbit anti-mCherry (1:500; Abcam, ab167453), Mouse anti-NompC (1:250; Liang et al. 2011), Rabbit anti-Sas4 (1: 350; gift from Jordan Raff), Rabbit anti-Dnah5 (1:2000; (zur Lage et al., 2021). The secondary antibodies (Life Technologies) were all used at a 1:500 concentration and included: Donkey anti goat (Alexa Fluor® 488, A11055), Donkey anti mouse (Alexa Fluor® 568, A10037), Donkey anti rabbit (Alexa Fluor® 647, A31573). Phalloidin (Life Technologies, A12380) was used at 1:2000.

### Confocal imaging

Images were captured using Leica TCS SP2, Nikon A1R and Zeiss Pascal confocal microscopes. Where appropriate, the length of protein staining of 4-6 randomly selected cilia in each z-stack was measured using Fiji software.

### Negative geotaxis climbing assay

10-15 mated female flies were collected for each group and placed in separate vials at 25°C for 24h to allow recovery. For each assay, flies from one group were placed in a 100-ml measuring cylinder, which was divided lengthwise into four 5-cm quadrants, and then after a 1-min recovery period, they were banged down to the bottom of the cylinder. Climbing performance in 10 s was recorded. The average climbing index was calculated based on the distance the flies climbed. The quadrant furthest away from the bottom is given a weight of 4 followed by 3, 2 and 1 to the quadrant nearest to the bottom and the climbing index = Σ (the numbers of flies in each quadrant) X (each quadrant weight) / the total number of flies. For comparing average climbing index, we conducted climbing assay for 5-7 times for each condition (each time with new flies), then calculated average climbing index.

### Statistical analysis

GraphPad Prism 9 was used for all analysis.

## RESULTS

To analyse protein localisation, we made extensive use of transgenic lines expressing the gene of interest (upstream, promoter and coding regions) fused to GFP, mVenus or mCherry reporters. In common with other motile ciliary genes in *Drosophila,* a relatively small proportion of upstream region drives chordotonal neuron expression through regulation by Rfx and Fd3F transcription factors (Newton et al., 2012). We focussed on the differentiating chordotonal neurons of Johnston’s Organ within the pupal antenna.

### TRP channels show differences in timing of localisation within chordotonal neuron cilia

Late in pupal development (96 h after puparium formation, APF), the chordotonal neuron cilium appears largely to have its mature structure. It is about 12-15 μm long, with proteins confined to proximal and distal zones separated by a ciliary dilation (Fig. 1). Most genetic studies of ciliary protein localisation examine adult or late-stage pupal neurons, and thereby restricting analysis to the final outcome of any transport or localisation process. In order to determine how localisation is achieved, we began by examining the time-course of localisation during chordotonal neuron ciliogenesis in the pupal antenna. At 6 h APF (shortly after the neurons are born), the cilium has already extended to c.4 μm (as estimated by the axonemal marker CG6652-GFP (Vieillard et al., 2016)) (Fig. 2A,B). Over the next few days, it extends slowly to its final length, as estimated by measuring the distance between Sas-4 (basal body marker) and NompC (distal marker, see later) (Fig. 2F–H). As the cilium grows exclusively from its tip, we deduce that the future proximal zone is generated in the first 24 h, and then further extension between 24 and 72 h adds the future distal zone. Extension appears to be coordinated with, and perhaps limited by, the elongation of the enclosing actin-rich scolopale and cap structures generated by support cells (Fig. 1C).

**Figure 2.**
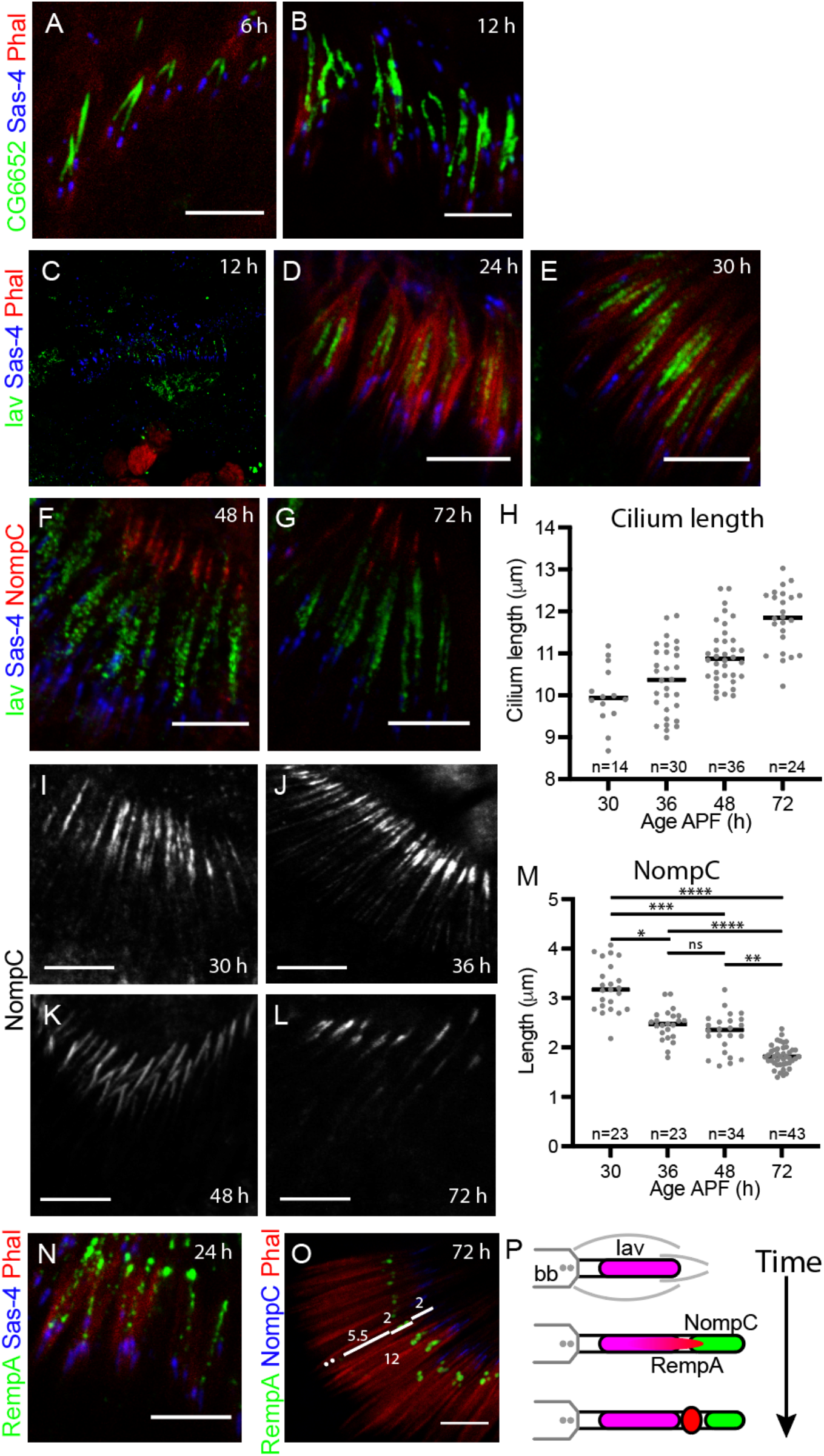
TRP channels localise to the cilium at different times. (A,B) The chordotonal cilia in pupal antenna show extension from at least 6 h APF onwards. Axoneme is marked with CG6652-GFP (green), basal bodies with Sas-4 (blue), F-actin-rich scolopales with phalloidin (red). (C–E) Iav-GFP localisation in early cilia. Iav-GFP (green), Sas-4 (blue), phalloidin (red). (C) At 12 h, Iav is not yet clearly within the cilia. (D) At 24 h, Iav shows substantial localisation to the proximal zone. (E) At 30 h, Iav localisation appears largely complete. (F,G) Iav-GFP localisation in later cilia. Iav-GFP (green), Sas-4 (blue), NompC (red). At 48 h (F) and 72 h (G), Iav-GFP and NompC are separated in the proximal and distal zones respectively. (H) Graph showing cilium length over time. Estimate based on measuring the distance between the basal body and tip of NompC staining. (I–L) Progression of NompC localisation. NompC shows localisation to the distal zone at 30 h (I), and this becomes more restricted over time (J–L). (M) Graph showing the restriction of NompC localisation to the distal zone over time. (N,O) RempA localisation to the ciliary dilation is progressive. (N) At 24 h, RempA is scattered in the cilium. RempA-YFP (green), Sas-4 (blue), Phalloidin (red). (O) At 72 h, RempA is confined to the ciliary dilation. RempA-YFP (green), NompC (blue), Phalloidin (red). (P) Schematic summary of ciliary protein localisation during ciliogenesis. All scale bars are 5 μm.

We examined the localisation of the two TRP channels Iav and NompC (Fig. 2C–L). Neither channel was detected in the first 6 h of ciliogenesis. TRPV subunit, Iav-GFP, was first observed at around 6 h but not clearly localised (Fig. 2C). But at 24 h, Iav-GFP was clearly localised along the growing length of the cilium from 2.5-3.5 μm to a maximum extent of 5-6 μm at 30-36 h (Fig. 2D,E). At later stages, the extent of staining appeared constant despite further cilium extension (Fig. 2F,G). This suggests that Iav localises directly to its proximal location as the proximal zone is being generated. In contrast, the distal zone channel NompC, was not detectable in the cilium until about 30 h. Upon its initial appearance, it localised directly to a 3-μm distal part of the cilium, with clear separation from Iav-GFP (Fig. 2F,G). Over the next few days as the tip extends, NompC remains in the distal part, and becomes restricted to a smaller area of the ciliary tip (c.1.8 μm) (Fig. 2I-M).

We examined localisation of ciliary dilation proteins to determine when this structure may become important in separating proximal and distal zones. Interestingly, RempA-YFP does not become fully localised to the future ciliary dilation until about 72 h (Fig. 2N,O). This suggests that the initial restriction of TRP proteins is not dependent on the ciliary dilation. To test whether expression timing determines TRP localisation, overexpressed NompC using a Gal4 driver that is active in chordotonal neurons from their birth *(scaGal4, UAS-NompC).* In these antennae, the localisation to the tip was more extended than normal, but NompC remained in the distal zone (Fig. S1). This suggests that timing of expression of NompC does not explain its exclusion from the proximal zone.

In summary, the time-course of TRP channel entry into the cilium correlates with their sites of localisation – proximal-early (Iav) and distal-late (NompC) (Fig. 2P).

### IDA motor complexes are generally distributed in the cilium until late in ciliogenesis

Although the role of DNAAFs in cytoplasmic pre-assembly within chordotonal neurons has been studied (Diggle et al., 2014; Moore et al., 2013; zur Lage et al., 2018, 2021), the later stages of dynein motor complex assembly have not been addressed previously. To follow IDA complex localisation, we used an mVenus fusion reporter for Dnali1, hereafter referred to as **IDA-Dnali1** for clarity (Moore et al., 2013). This is a predicted light-intermediate chain subunit of several monomeric IDA species (zur Lage et al., 2019) (Table 1). The corresponding *Chlamydomonas* subunit (p28) is required for cytoplasmic IDA complex pre-assembly and so IDA complexes do not localise in its absence. In the adult antenna, we found that IDA-Dnali1 localised to approximately 4.5-5.5 μm structures, corresponding well to the extent of the proximal zone as observed by both transmission electron microscopy and Iav localisation (Fig. 1; Fig. 3A).

**Figure 3.**
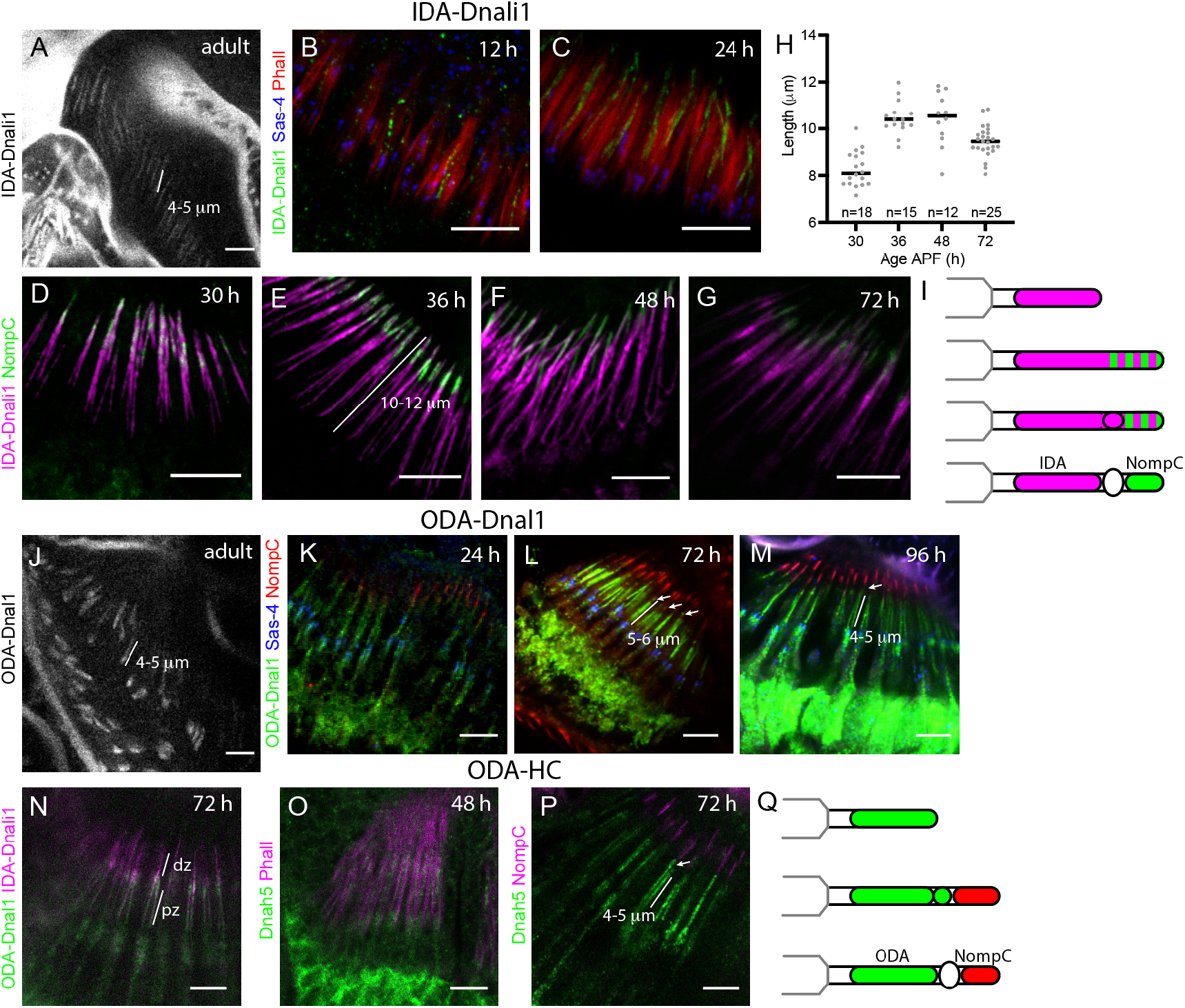
ODA and IDA markers localise differently during ciliogenesis. (A–I) Localisation of IDA-Dnali1-mVenus. Pupal antennae at times indicated. (A) Adult antenna (mVenus fluorescence). (B,C) IDA-Dnali1 enters the cilium early. IDA-Dnali1 (green), Sas-4 (blue), Phalloidin (red). (D–G) IDA-Dnali1 localisation extends into the NompC-containing distal zone. IDA-Dnali1 (magenta), NompC (green). (H) Graph showing length of Dnali1-mVenus staining. For each time point. (I) Schematic summary of IDA-Dnali1 localisation. (J–Q) Localisation of ODA-Dnal1-mVenus. Pupal antennae at times indicated. (J) Adult antenna (mVenus fluorescence). (K–M) Pupal antennae, ODA-Dnal1 (green), Sas-4 (blue), NompC (red). (K) ODA-Dnal1 enters the cilium early. (L) At 72 h, ODA-Dnal1 localisation is confined to the proximal zone and an accumulation at the ciliary dilation (arrows) (M) At 96 h, ODA-Dnal1 localisation is confined to the proximal zone only (arrow marks the location of ciliary dilation). (N) 72-h pupal antenna, showing location of ODA-Dnal1-mVenus (green) and IDA-Dnali1-mCherry (magenta). Both are in the proximal zone (pz) but only IDA-Dnali1 extends into the distal zone (dz). (O,P) Anti-Dnah5 labelling. (O) At 48 h, Dnah5 is present in the proximal zone. Dnah5 (green), phalloidin (magenta). (P) At 72 h, Dnah5 is in the proximal zone and ciliary dilation (arrow). Dnah5 (green), NompC (magenta). (Q) Schematic summary of ODA marker localisation. All scale bars are 5 μm.

**Table 1.** Genes/proteins in this study.

| *Drosophila* |  | Human | *Chlamydomonas* | Notes |
| --- | --- | --- | --- | --- |
| *iav* | CG4536 | TRPV1-6 |  | TRP channel |
| *nompC* | CG11020 | *trpn1* (zebrafish) |  | TRP channel |
| *ODA-Dnal1* | CG8800 | *DNAL1* | LC1 | ODA light chain |
| *IDA-Dnali1* | CG6971 | *DNALI1* | p28 | IDA light-intermediate chain |
| *Dnah5* | CG9492 | *DNAH5* | ODA2 | ODA dynein heavy chain (gamma) |
| *Dhc93AB* | CG3723 | *DNAH11* | ODA4 | ODA dynein heavy chain (beta) |
| *Ccdc114* | CG14905 | *CCDC114* | DC2 | ODA docking complex |
| *Ccdc151* | CG14127 | *CCDC151* | ODA10 | ODA docking complex |
| *Ccdc39* | CG17387 | *CCDC39* | FAP59 | Molecular ruler |
| *Cfap57* | CG4329 | *CFAP57/WDR65* | FAP57 | IDA docking protein |
| *rempA* | CG11838 | *IFT140* | IFT140 | IFT-A complex |
| *dTulp/ktub* | CG9398 | *TUB, TULP1, TULP3* | TLP1 | TRP transport adaptor |

Like Iav, IDA-Dnali1 enters the cilium early in ciliogenesis (at least by 12 h, Fig 3B,C). In contrast to Iav, however, IDA-Dnali1 it is detected along the length of the cilium as it extends further, to a maximum of 10.5 μm at 48 h (Fig. 3H). Indeed, there is substantial overlap with NompC at 30-48 h (Fig. 3D-F), suggesting that IDA-Dnali1 is present in both proximal and distal zones. From 72 h, IDA-Dnali1 becomes progressively refined to the proximal zone (9 μm at 72 h) (Fig. 3G), but this refinement is still not complete by 96 h APF. Overall, this suggests that IDA-Dnali1, and therefore IDA complexes, are present along the entire cilium during ciliogenesis until a late stage of maturation, when they become confined to the proximal zone (Fig. 3I).

### ODA motor complexes are distributed in proximal zone and ciliary dilation until late in ciliogenesis

A different pattern was observed for the ODAs. To follow ODA complex localisation, we used a reporter for the light chain subunit CG8800 (homologue of human DNAL1 and *Chlamydomonas* LC1), hereafter referred to as **ODA-Dnal1** (zur Lage et al., 2019). In the adult antenna, ODA-Dnal1 localised to approximately 4.5-5.5 μm structures corresponding to the proximal zone (Fig. 1; Fig. 3J). Like IDA-Dnali1, ODA-Dnal1 is present in the cilium early but differs in that it is excluded from the distal cilium as it later forms, thereby showing no overlap with NompC at 30-96 h (Fig. 3K-M). However, compared with Iav, the extent of ODA-Dnal1 localisation appears longer at 72 h (c.6 μm), suggesting that it extends into the future ciliary dilation (Fig. 3L). By 96 h, the extent of localisation reduces to resemble that of Iav (c.5 μm), thereby becoming restricted to the proximal zone (Fig. 3M). The difference between IDA-Dnali1 and ODA-Dnal1 localisation was confirmed by simultaneous detection of IDA-Dnali1-mVenus and ODA-Dnal1-mCherry (Fig. 3N). To confirm that the localisation dynamics of ODA-Dnal1 reflected ODA complexes, we used an antibody generated against *Drosophila* ODA HC, Dnah5 (CG9492). Dnah5 was localised in a similar manner to ODA-Dnal1 (Fig. 3O,P).

Therefore, we infer that proximal zone proteins (dynein complexes and Iav) all enter the cilium early, but there are differences in initial localisation. Iav never localises beyond the proximal zone. ODA-Dnal1 and Dnah5 appear similar to Iav in never being present at the future distal zone, but they transiently extend into the area of the ciliary dilation (Fig. 3Q). In contrast, IDA-Dnali1 is generally distributed throughout the cilium, only becoming confined to the proximal region late in cilium maturation (Fig. 3I).

Since TRPV and dynein complexes ultimately colocalise at the proximal zone, and are functionally connected (Karak et al., 2015), we asked whether their localisation was co-dependent. We first analysed flies with a CRISPR-engineered null mutation of *Dnaaf3 (CG17669),* the orthologue of the human dynein assembly factor and PCD gene, *DNAAF3.* Human *DNAAF3* is required for all dynein assembly, and *Drosophila Dnaaf3^CR^* mutant flies show absence of dynein arms from the chordotonal neuron cilia (zur Lage et al., 2021). However, we found that Iav localisation was unaltered in *Dnaaf3^CR^* mutant antennae (Fig. 4A,B). Conversely, IDA-Dnali1 and ODA-Dnal1 were unaltered in dTulp1 knockdown antennae *(scaGal4,* UAS-*dTulp*-RNAi) (Fig. 4C–E), in which Iav does not enter the cilium (Fig. 4F,G) (Park et al., 2013). This suggests that dynein transport and localisation require neither Iav nor the dTulp/PIP mechanism of ciliary targeting.

**Figure 4.**
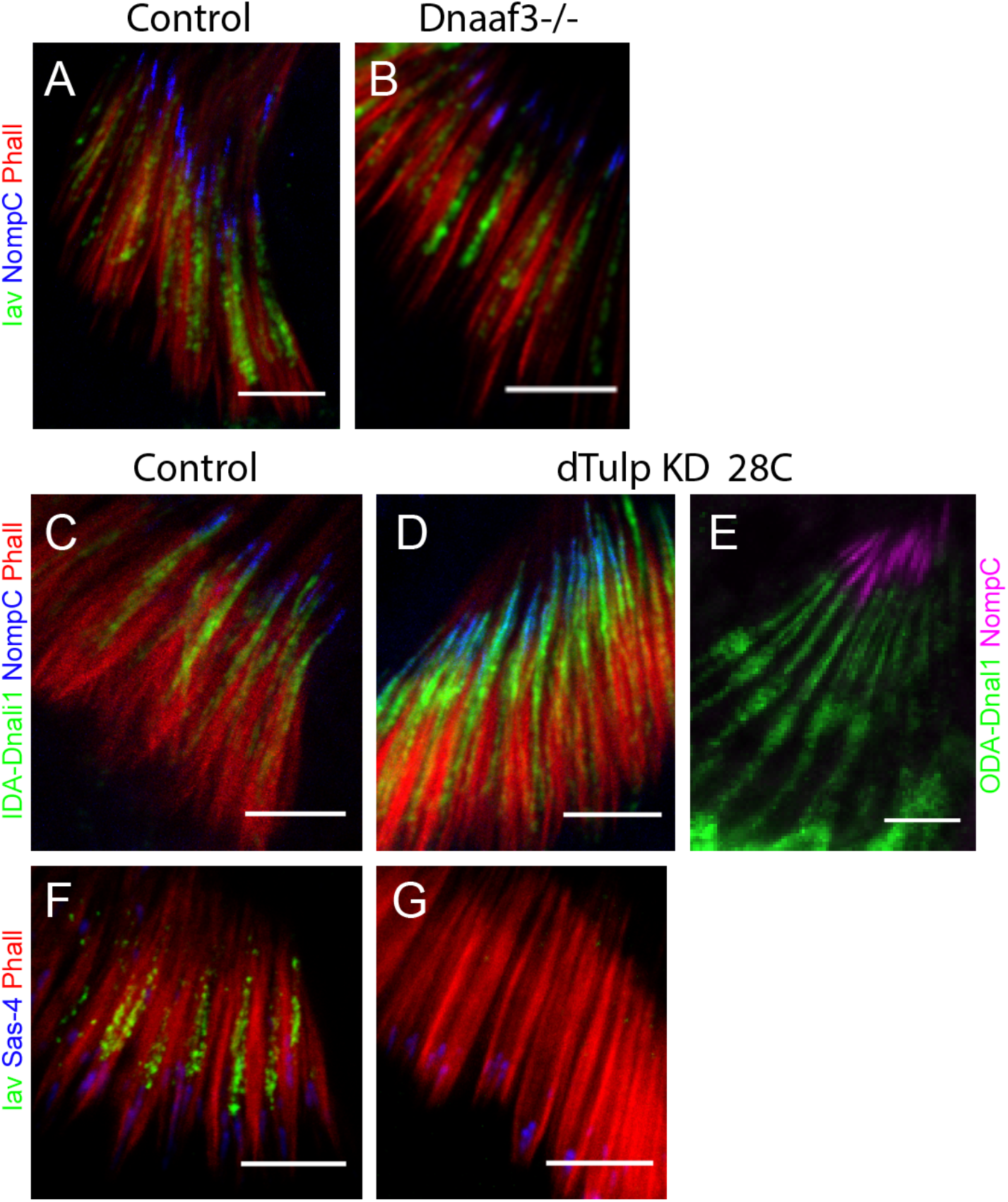
Iav and IDA-Dnali1 localise to the cilium independently of each other. Pupal antenna at approximately 72 h. (A,B) Loss of dynein complexes from the cilium *(Dnaaf3* homozygotes) does not prevent localisation of Iav. Iav (green), NompC (blue), phalloidin (red). (A) *Dnaaf3* heterozygote control. (B) *Dnaaf3* homozygote. (C,D) Knockdown of dTulp does not prevent localisation of IDA-Dnali1. IDA-Dnali1 (green), NompC (blue), phalloidin (red). (C) Control *(scaGal4/+).* (D) dTulp knockdown *(scaGal4/UAS-dTulp-RNAi.* (E) dTulp knockdown *(scaGal4/UAS-dTulp-RNAi)* does not prevent ODA-Dnal1 localisation. ODA-Dnal1 (green), NompC (blue), phalloidin (red). Note that NompC localisation spreads proximally upon dTulp knockdown as previously reported (Park et al., 2013), but still does not overlap with ODA-Dnal1, suggesting NompC remains confined to the distal zone. (F,G) Confirmation that Iav does not enter the cilium upon dTulp knockdown (Park et al., 2013). Iav (green), Sas-4 (blue), phalloidin (red). Iav localises in the control (F) but not in knockdown antenna (G). All scale bars are 5 μm.

### IDA docking proteins are generally distributed in the cilium until late in ciliogenesis

To understand further how dynein complexes populate the proximal zone, we analysed *Drosophila* orthologues of proteins required for dynein docking to the axoneme. In *Chlamydomonas,* the binding of IDA complexes in their correct periodicity requires two coiled-coil proteins that are said to form a ‘96 nm molecular ruler’ (Fig. 1A; Table 1). The human homologues of these, CCDC39 and CCDC40, cause PCD when mutated, with disruption of binding of IDA complexes, nexin-dynein regulatory complexes (N-DRC) and radial spoke complexes (Antony et al., 2013; Becker-Heck et al., 2011; Merveille et al., 2011). *Drosophila* has a single orthologue of each gene: *CG17387/Ccdc39* and *l(2)41Ab/Ccdc40* (zur Lage et al., 2019). To confirm that *Drosophila Ccdc39* is required for IDA localisation, we carried out RNA interference. Knockdown of *Ccdc39* in chordotonal neurons *(scaGal4,* UAS-*Ccdc39*-RNAi) resulted in reduced IDA-Dnali1 localisation in the cilium but localisation of ODA-Dnal1 appeared unaffected, consistent with a role in stable docking of IDAs (Fig. S2). Residual IDA-Dnali1 protein in the cilium may represent complexes that enter the cilium but are not stably docked.

We examined the localisation of a Ccdc39-mVenus fusion protein, and observed that it was present along the whole length of the developing cilium from 24 h to 96 h (Fig. 5A,B). This was confirmed by finding the complete colocalization of Ccdc39-mVenus with IDA-Dnali1-mCherry (Fig. 5C). Therefore, like IDA-Dnali1 itself, an IDA docking protein is not restricted to the proximal zone until late in ciliogenesis (Fig. 5E). Another protein required for a subset of IDA motors (g and d) is CFAP57 (Bustamante-Marin et al., 2020; Lin et al., 2019), and mutants of the *Drosophila* orthologue gene *CG4329,* are moderately deaf (Senthilan et al., 2012). In contrast to Ccdc39, we found that *Cfap57-* GFP (Augière et al., 2019) shows greater exclusion from the distal zone (Fig. 5D).

**Figure 5.**
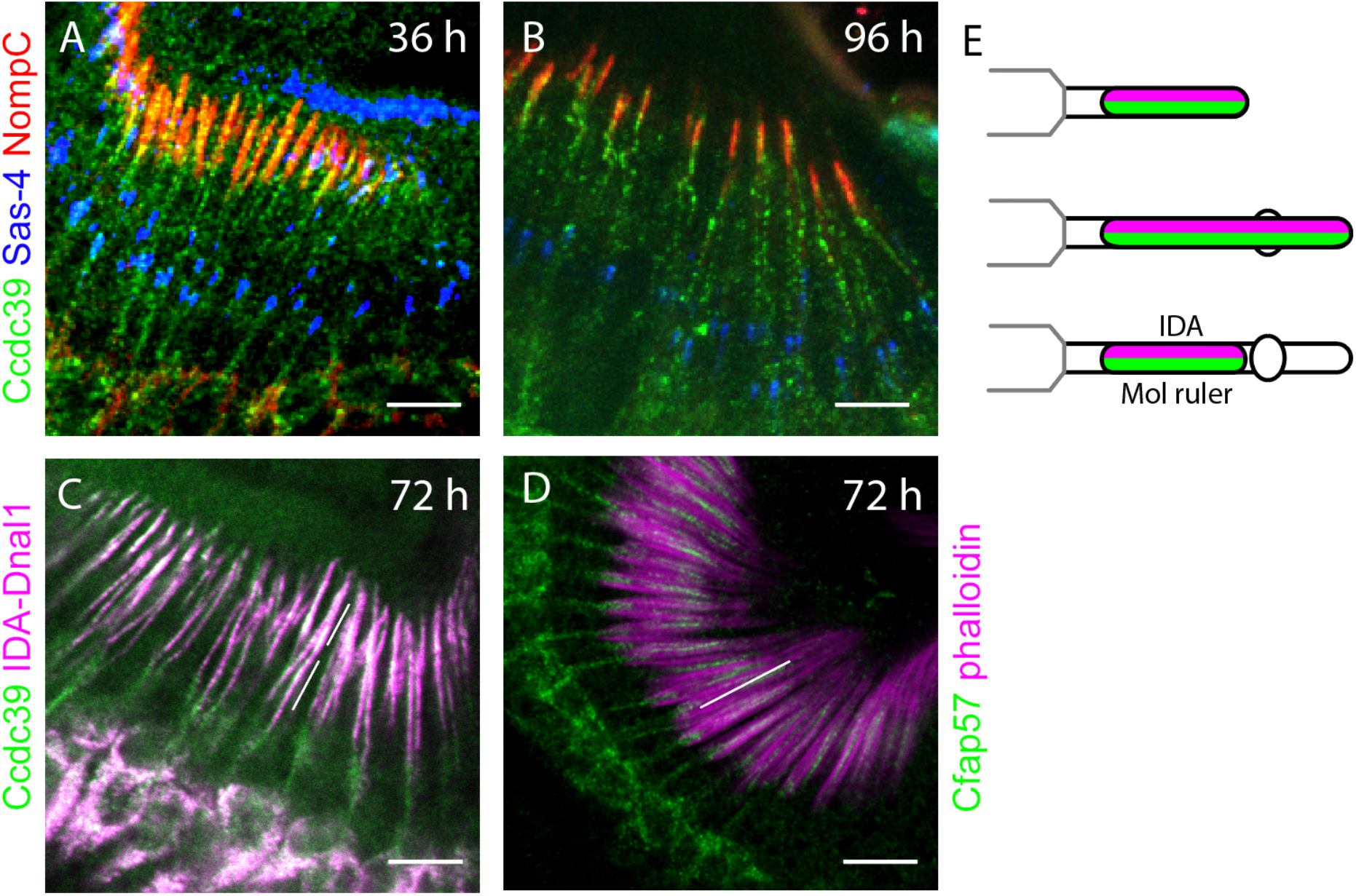
Ccdc39-mVenus localises to the whole cilium in pupal antennae. (A,B) Immunofluorescence for Ccdc39-mVenus (green), Sas-4 (blue), NompC (red). (A) 36-h pupal antenna. (B) 96-h pupal antenna. (C) 72-h pupal antenna. Colocalisation of Ccdc39-mVenus (green) and IDA-Dnali1-mCherry (magenta). (D) Localisation of Cfa57-GFP (green) and phalloidin (magenta). (E) Schematic summary of localisation. All scale bars are 5 μm.

### An ODA docking complex (ODA-DC) subunit, *Ccdc114,* is required for stable docking but not cilium entry of ODAs

In order to characterise stable ODA localisation, we wished to analyse the function of *Drosophila* ODA-DC. Although there are differences between *Chlamydomonas* and human ODA-DC (Desai et al., 2018), a common component is the coiled-coil protein known as DC2 in *Chlamydomonas (oda1* gene), and conserved in humans as CCDC114. In *Drosophila,* an *oda1/CCDC114* homologue, *CG14905,* is expressed exclusively in chordotonal neurons (Jerber et al., 2014; zur Lage et al., 2019) (Table 1). The following experiments present evidence that this gene is a functional homologue of *DC2/CCDC114,* and hereafter we refer to the *CG14905* gene as *Ccdc114.*

A null allele of *Ccdc114* was generated using CRISPR/Cas9 genome engineering to replace the ORF with a mini-*white* gene. Homozygous *Ccdc114^CR^* flies were viable and fertile, but exhibited reduced ability in a climbing assay, which (combined with the gene’s chordotonal neuron-specific expression pattern) is consistent with defective chordotonal neuron function (Fig. S3A). This phenotype could be partially rescued by an mVenus-tagged *Ccdc114* gene expressed under its own promoter (Fig. S3A).

We determined the requirement of *Ccdc114* for ODA localisation. At 36, 72 and 96 h, ODA-Dnal1 and Dnah5 localisation were both largely absent in *Ccdc114^CR^* mutant cilia, suggesting that the anchoring of ODA complexes in the proximal zone was abolished (Fig. 6A-F). This supports the prediction that *Ccdc114* encodes a subunit of the ODA-DC that is essential for stable ODA docking. In the mutant cilia at 36 and 72 h, however, some remaining ODA protein (ODA-DNAL1 and Dnah5) was detected in the region of the ciliary dilation. This accumulation disappeared at 96 h (Fig. 6E,F). This supports the hypothesis suggested above that there are separate pools of ODAs within differentiating cilia: stable/ODA-DC-dependent in the proximal zone and transient/ODA-DC-independent at the ciliary dilation. Moreover, the observations demonstrate that in the absence of ODA-DC, ODA motor complexes are still able to enter the cilium independently (Fig. 6K). Together, these results confirm that Ccdc114, and by inference ODA-DC, is required for stable localisation of ODA complexes in the proximal zone.

**Figure 6.**
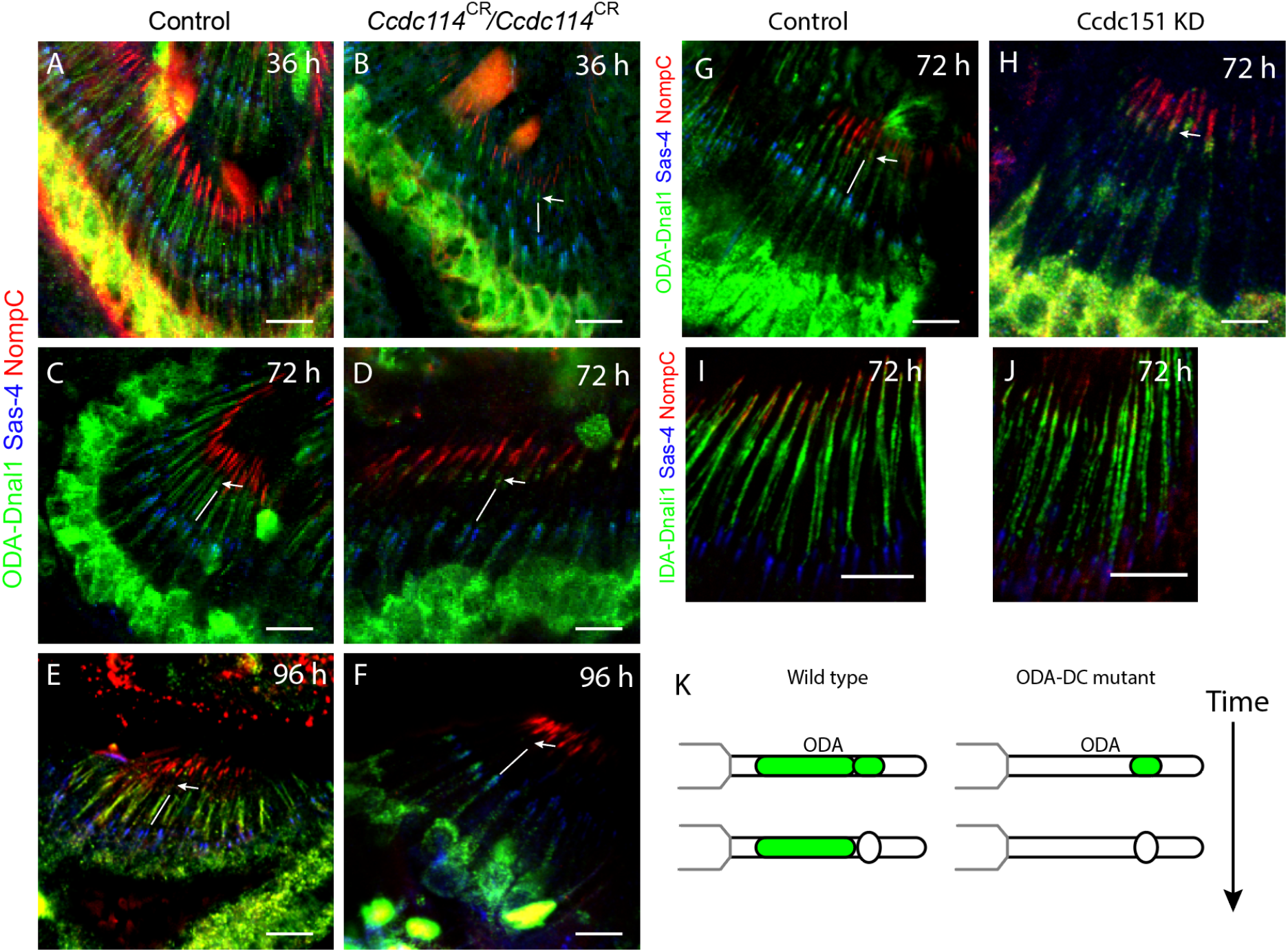
Loss of stable ODA localisation in ODA-DC mutants. (A-F) Localisation of ODA-Dnal1 to the proximal zone is abolished in Ccdc114 homozygotes (B,D,F) compared with heterozygote controls (A,C,E) in pupal antennae of the timepoints indicated. ODA-Dnal1 (green), Sas-4 (blue), NompC (red). At 36 and 72 h, expression is absent from the proximal zone (white line) but remains in the ciliary dilation (arrow). At 96 h, all expression is absent. (G,H) Localisation of ODA-Dnal1 to the proximal zone (white line) is strongly reduced in knockdown of Ccdc151 (scaGal4, UAS-Ccdc151-RNAi) (G) compared with control (scaGal4, KK line control) (H). As for Ccdc114, expression remains in the ciliary dilation (arrow). ODA-Dnal1 (green), Sas-4 (blue), NompC (red). In contrast, localisation of IDA-Dnali1 to cilia is unaffected in knockdown of Ccdc151. IDA-Dnali1 (green), Sas-4 (blue), NompC (red). (K) Schematic summary. All scale bars are 5 μm.

To confirm this finding, we examined the function of a second predicted ODA-DC component, *CG14127,* a homologue of *ODA10/CCDC151* (Table 1). Although the *Chlamydomonas ODA10* gene is associated with ODA trafficking rather than docking (Desai et al., 2018; Fowkes and Mitchell, 1998), vertebrate *CCDC151* appears to be a component of ODA-DC itself (Gui et al., 2021; Hjeij et al., 2014; Jerber et al., 2014). In order to investigate whether *CG14127* (hereafter *Ccdc151)* affects dynein complex localisation in *Drosophila,* we used genetically supplied RNAi interference. *Ccdc151* knockdown in sensory neurons *(scaGal4,* UAS-*Ccdc151*-RNAi) resulted in flies with sensory defects, consistent with defective chordotonal neuron function (Jerber et al., 2014; zur Lage et al., 2019) (Fig. S3B). Knockdown resulted in strong loss of ODA-DNAL1 protein from the proximal zone, but the transient protein accumulation at the ciliary dilation remained, similar to the phenotype observed in the *Ccdc114* mutant (Fig. 6G,H). A previous study suggested that zebrafish *ccdc151* may be required for IDA localisation (Jerber et al., 2014). In *Drosophila,* we found that IDA-Dnali1 localisation to the axoneme appeared unaffected by *Ccdc151* knock-down (Fig. 6I,J). Together, these results are consistent with *Ccdc151* being a component of ODA-DC.

### Ccdc114 protein is targeted directly to the proximal zone

To determine the localisation time course of ODA-DC, we examined the ciliary localisation of *Ccdc114-mVenus* fusion protein. From 24 h onwards, the fusion protein localised to the proximal zone of the chordotonal neuron cilia, consistent with its docking function (Fig. 7A-C). During ciliogenesis, the extent of Ccdc114-mVenus localisation at no time exceeded 5 μm, which is similar to Iav-GFP. Thus, Ccdc114 seems to be targeted specifically and directly to the future proximal zone, in contrast to Ccdc39. Notably, the extent of Ccdc114 localisation is shorter than we had observed for ODA-Dnal1, with the former not extending into the ciliary dilation area. This difference was confirmed by co-labelling of Ccdc114-mVenus and ODA-Dnal1-mCherry (Fig. 7D,I). These observations are consistent with the hypothesis that ODAs transiently accumulate independently of ODA-DC at or near the future ciliary dilation.

**Fig. 7.**
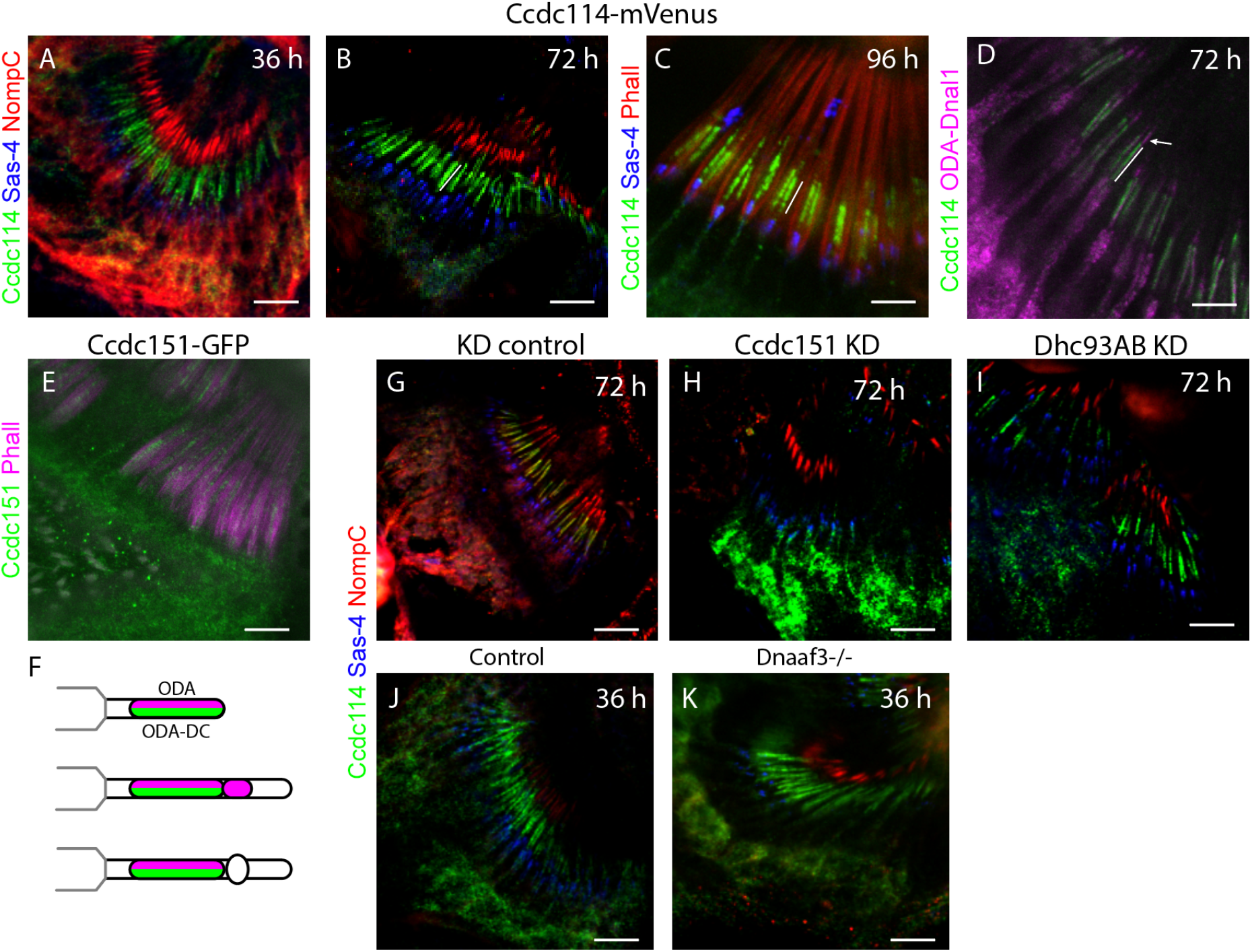
ODA-DC components localise directly to the proximal zone. (A-C) Pupal antennae at indicated hours, showing localisation of Ccdc114-mVenus to the proximal zone (white line). Ccdc114-mVenus (green), Sas-4 (blue), NompC (red, A,B) or phalloidin (red, C). (D) Colocalisation of Ccdc-115-mVenus (green) and ODA-Dnal1-mCherry (magenta) confirms colocalization in the proximal zone (line) but only ODA-Dnal1-mCherry localises beyond (arrow). (E) Ccdc151-GFP (green) localises to cilia at 72 h; phalloidin (red). (F) Schematic summary of ODA-DC and ODA localisation. (G-K) Pupal antenna showing Ccdc151 (green), Sas-4 (blue), NompC (red). (G,H) Knockdown of Ccdc151 *(scaGal4,* UAS-*Ccdc151*-RNAi) results in loss of Ccdc114 from the cilia (H) compared with control *(scaGal4,* KK control) (G). (I) Knockdown of Dhc93AB *(scaGal4,* UAS-*Dhc93AB-RNAi)* does not affect Ccdc114 in the cilia. (J,K) Ccdc114 is localised normally in Dnaaf3 homozygous mutant antennae (K) compared with control (J). All scale bars are 5 μm.

**Figure 9.**
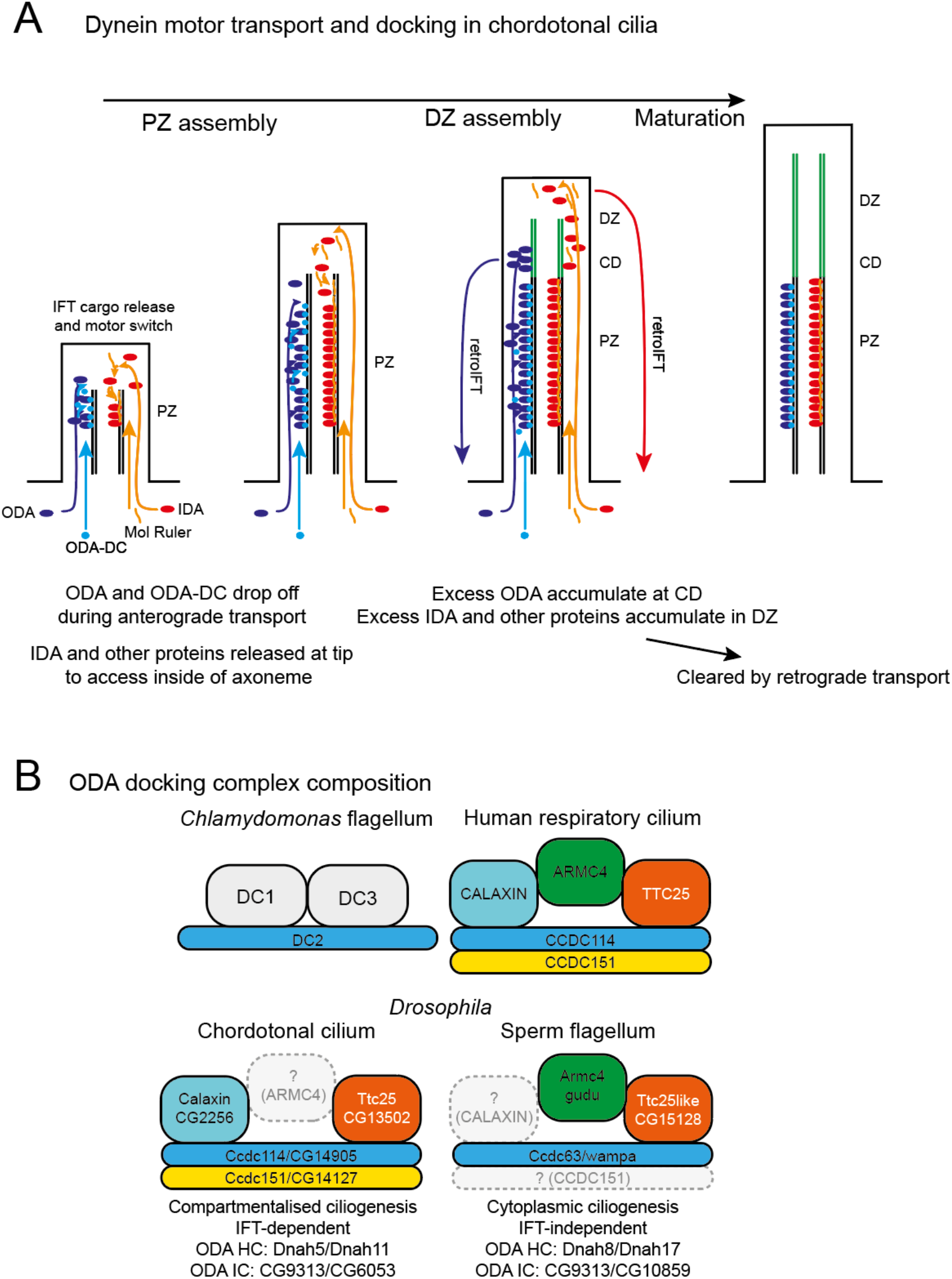
Models of dynein transport in chordotonal ciliogenesis and of ODA-DC composition. (A) ODA and ODA-DC are able to access their microtubule binding sites along the length of the proximal zone (PZ). They are released from IFT trains before they reach the ciliary tip. Consequently, as the distal zone (DZ) extends, excess complexes are not found far beyond the proximal zone – transient accumulation occurs at the future ciliary dilation (CD). In contrast, IDA, their docking proteins, and other components access their microtubule binding sites at the growing tip only, and are released from IFT trains at the tip. Excess complexes accumulate transiently in the distal zone. In all cases, we envisage that excess complexes (not stably bound to the PZ) are eventually cleared by retrograde IFT. (B) Schematic of the proposed composition of *Drosophila* ODA-DC complexes, relative to the counterparts in *Chlamydomonas* flagella and human respiratory cilia. Colour coding indicates homology. Expression and functional evidence strongly suggest the presence of substantially different ODA-DCs in chordotonal cilia compared with sperm flagella. These differences may be related to differences in mode of ciliogenesis or in differences in ODA composition.

Analysis of Ccdc151-GFP localisation showed a similar pattern of localisation to Ccdc114 (Fig. 7E), confirming that Ccdc151 and Ccdc114 are likely physically interacting as part of ODA-DC (Hjeij et al., 2014). Given this, we examined the dependence of Ccdc114 localisation on Ccdc151 function, with the finding that upon Ccdc151 knockdown, Ccdc114 protein was completely absent from cilia (Fig. 7F,G). Similar observations were reported for zebrafish morphants and human PCD patients (Hjeij et al., 2014; Jerber et al., 2014; Onoufriadis et al., 2013), suggesting that Ccdc151 is a core component of the *Drosophila* ODA-DC.

We then examined Ccdc114 protein localisation in flies in which ODA complexes are not formed. We examined antennae in which the expression of ODA heavy chain *Dhc93AB* is knocked down in chordotonal neurons *(sca-Gal4; UAS-Dhc93AB RNAi). Dhc93AB* encodes a *DNAH11* orthologue (Table 1) that is known to be expressed and required in chordotonal neurons (zur Lage et al., 2019). In confirmation of this, RNAi knockdown of *Dhc93AB* resulted in flies with reduced climbing ability (Fig. S3B). The reduction of the heavy chain is predicted to result in failure of ODA motor pre-assembly and therefore failure in localisation of the ODA complex (King, 2016). Upon *Dhc93AB* knockdown, however, localisation of Ccdc114-mVenus appeared unaltered (Fig. 7F,H). These results suggest that ODA-DC is transported and bound to the proximal zone axoneme independently of ODA motor complexes. To extend this observation, we analysed localisation in *Dnaaf3^CR^* mutant flies. In homozygous *Dnaaf3^CR^* mutant antennae, Ccdc114-mVenus localisation appeared little different from wildtype (Fig. 7B). Together, these results suggest that ODA-DC is transported and bound to the proximal zone axoneme independently of ODA motor complexes.

## DISCUSSION

In this study, we characterise the dynamics of protein localisation to zones within the highly sub-compartmentalised chordotonal neuron cilium. Ciliogenesis in the chordotonal neurons of Johnston’s Organ is prolonged, occurring over several days of pupal development. Over this time, TRP channel proteins appear to be targeted directly to their respective zones, albeit at different times. In contrast ODAs and IDAs are initially not confined to the proximal zone until late into ciliary maturation. ODA and IDA markers transiently accumulate at the ciliary dilation and whole distal region, respectively (Fig. 8A). Below we discuss the roles of docking factors in dynein motor targeting, and possible implications and mechanisms.

### Spatial differences in TRP channel localisation correlate with temporal differences in cilium entry

During transport, ciliary TRP channels are linked to IFT trains by Tulp adaptors, which release their cargoes in response to PIP levels within the cilium. For *Drosophila* chordotonal cilia, dTulp1 is required for correct targeting of both Iav and NompC (Park et al., 2013), and this is affected by PIP signalling regulated by *dInpp5e* (Park et al., 2015). It is not clear, however, how the different localisations of the two TRP channels are achieved. Perhaps in response to PIP levels, Iav is released from IFT trains sooner than is NompC. In this possibility, differences in drop-off from IFT trains (processivity) may provide the initial localisation. Our observations suggest instead that Iav enters the cilium early in ciliogenesis and populates the proximal zone as it is being generated, while NompC enters later and populates the distal zone as it is being generated subsequently. The time of entry into the cilium suggests a simple mechanism for TRP protein localisation to different zones based on timing of expression, but we found that early overexpression of NompC did not cause substantial mislocalisation to the proximal zone. It remains possible that timing of entry into the cilium could be controlled, rather than timing of expression.

Since the zones of TRP channel localisation are separated by the ciliary dilation, it has been proposed that this structure actively defines the zones and targets proteins to them. Indeed, disruption of the proteins that locate to, and are required for, the ciliary dilation also affect Iav and NompC localisation. However, we find that ciliary dilation proteins are not localised until quite late, whereas Iav and NompC are largely targeted to their respective zones immediately upon ciliary entry. It seems more likely therefore that the role of the ciliary dilation may be in later refinement and maintenance of this initial separation. It remains unclear, therefore, what determines the initial localisations of the TRP channels. Since the cilium is ensheathed in scolopale and cap structures from an early stage, one plausible mechanism is that the surrounding cap provides an external cue for distal proteins such as NompC, e.g. via contact between the cap and ciliary tip membrane mediated by the linking protein, NompA (Chung et al., 2001). Dynamic changes in such contact during maturation might also explain why the extent of NompC localisation becomes more restricted at later stages of development. A second possibility is a temporal change in axonemal modifications mechanisms during synthesis of the proximal and then distal zones.

### Dynein transport in chordotonal cilia: differences between ODAs and IDAs correspond to differences in IFT dynamics known in *Chlamydomonas*

Very little is known about the dynamics of dynein motor transport and localisation in metazoans. We find that ODA and IDA complexes enter the cilium early in ciliogenesis of chordotonal neurons. In addition to localisation in the proximal zone, both complexes show temporary accumulation beyond their final destination. For ODA, accumulation is confined to a point just beyond the proximal zone, whereas for IDA it is at the whole of the ciliary tip. We suggest that these represent transient populations of (excess?) motor complexes that are eventually cleared from the cilium, presumably by retrograde IFT.

Why the difference in non-proximal ODA and IDA populations? A possibility is that this reflects differences in how the complexes are released from IFT trains. This is based on comparison with observations obtained in *Chlamydomonas* from live imaging and dikaryon analysis (observing the assembly of complexes onto pre-existing flagella after dikaryon fusion). In these studies, most IFT cargoes are transported to the flagellar tip before release (Qin et al., 2004; Wren et al., 2013). Such a mechanism has been described for IDAs (Piperno et al., 1996; Viswanadha et al., 2014), radial spoke complexes (Johnson and Rosenbaum, 1992; Lechtreck et al., 2018), nexin-DRCs (Bower et al., 2013; Wren et al., 2013) and central pair components (Lechtreck et al., 2013). Release at the tip is thought to occur by ‘maturation’ and remodelling of IFT trains, probably an IFT-A function. Thus, most of these IFT cargoes show a high processivity (low rate of dissociation from IFT trains) during transport (Chien et al., 2017; Dai et al., 2018; Wren et al., 2013). The complexes are then presumed to diffuse locally onto docking sites accessible at the growing tip of the cilium. Thus, we speculate that the distal zone accumulation of IDA complexes represents continued transport to the tip throughout ciliary growth, but lack of local docking sites in the distal zone means that these complexes are eventually cleared.

It has been suggested that transport to the tip is required for IDAs (and many other components such as radial spokes, nexin-DRC) because their docking sites in the interior of the axonemal shaft are only accessible via the tip during ciliogenesis (Fig. 1B) (Ahmed et al., 2008; Piperno et al., 1996; Piperno and Mead, 1997; Qin et al., 2004). In contrast, ODA motor complexes can theoretically access their docking sites (when present) along the entire length of the axoneme. Thus, ODA may have low processivity (high rate of dissociation) and so may be released from IFT trains along the length of the cilium before diffusing locally onto docking sites (Dai et al., 2018). This may explain the lower level of transient accumulation beyond the proximal zone – most ODA complexes are released before the trains reach the distal zone. It is not known how the release might be regulated, but it does not appear to involve the Tulp/PIP mechanism.

Little is known of ODA-DC transport and localisation, and it has not yet been demonstrated that it is an IFT cargo, although several metazoan ODA-DC proteins are known to associate with IFT proteins (Wallmeier et al., 2016; Xu et al., 2015). In chordotonal cilia, ODA-DC (as marked by Ccdc114) is directly targeted to the proximal zone, independently of ODA complexes. This pattern of localisation suggests that ODA-DC is either a low processivity IFT cargo, like ODA motors, or that it populates the cilium via simple diffusion from the base (Owa et al., 2014).

To understand how dynein motors stably dock only in the proximal zone, it will be necessary to understand how their docking sites are eventually restricted to this zone. For the 96-nm molecular ruler subunit, Ccdc39, its early distribution along the whole cilium is consistent with it being released from IFT trains only at the tip. Why it does not bind to the distal zone (and thereby guide the docking of IDAs) is not presently clear. It seems highly probable that the axoneme has different properties in the proximal and distal zones that allow differential binding of docking proteins, and thereafter of dynein complexes. One possibility is that there is a difference in tubulin modification in the proximal and distal axonemes. Another is that there are initial differences in so-called ‘microtubule internal proteins’ (MIPs) (Ma et al., 2019; Owa et al., 2019). These are proteins that clearly can only incorporate within the microtubules at the growing tips. Their roles are only now being unravelled, but so far they appear to play an important role in defining the periodicity of binding of proteins to the outside of the microtubules. We predict that one or more key MIPs are incorporated only into the proximal zone during axoneme extension.

Overall, very little is known of compartmentation within cilia. Investigating dynein motor docking proteins in chordotonal cilia offer a useful model for exploring this further.

### Variation in the composition and function of metazoan ODA-DC

In the course of this investigation, we have begun to characterise the ODA-DC of chordotonal cilia. The composition of the ODA-DC has not been well characterized outside *Chlamydomonas,* although existing evidence indicates that the metazoan complex differs from that of the alga. Of the three *Chlamydomonas* ODA-DC subunits (DC1, DC2 and DC3), only DC2 is conserved as CCDC114 in vertebrates (Jerber et al., 2014; Onoufriadis et al., 2013) (Fig. 8B). We provide strong evidence that *CCDC114/DC2* is conserved in *Drosophila* as *CG14905/Ccdc114.* This gene is essential for ODA docking in chordotonal cilia. Moreover, it is clear that Ccdc114 is transported and localised independently of ODA complexes.

Until recently, the remaining proteins of the metazoan ODA-DC were not fully known. ODA-DC in human respiratory cilia is suggested to include *CCDC114, CCDC151,* and *ARMC4.* Mutation of these human genes results in PCD with loss of ODAs on the axoneme (Hjeij et al., 2013; Knowles et al., 2013; Onoufriadis et al., 2014). In humans, *TTC25* is required for ODA-DC localisation and function, but its axonemal localisation is retained in the absence of *CCDC114* function (Wallmeier et al., 2016), making it unclear whether it was part of the complex. Recent ultrastructural analysis, however, has shown that mammalian ODA-DC is a pentameric complex comprising the above subunits, TTC25 and a further subunit, Calaxin/EFCAB1 (Gui et al., 2021) (Fig. 8B).

Our evidence suggests that the subunits of this complex are conserved in *Drosophila* ODA-DC, but their requirement seems to vary between cell types. We found that Ccdc114 and Ccdc151 are likely part of ODA-DC in chordotonal cilia, but notably, neither are expressed in sperm and so cannot account for ODA docking in sperm flagella (zur Lage et al., 2019). In sperm, Ccdc114 expression is replaced by a paralogue, *wampa (CG17083)* (zur Lage et al., 2019). One feature of *wampa* mutant sperm is that their flagella lack ODAs (Bauerly et al., 2020), consistent with an ODA-DC function. Likewise, the *Drosophila TTC25* homologue, *CG13502 (Ttc25),* is restricted to chordotonal neurons, although it may be represented in sperm by a more distantly related gene, *CG15128 (Ttc25like).* The single *Drosophila ARMC4* homologue, *gudu,* is only expressed in sperm (zur Lage et al., 2019), where it is required for fertility (Cheng et al., 2013), whereas *Calaxin (CG2256)* is only expressed in chordotonal neurons. We therefore suggest that docking complex composition may differ substantially between chordotonal cilia and sperm flagella (Fig. 8B). Interestingly, this might be a common feature of metazoans: in humans *CCDC114* is required for respiratory cilia but PCD patients with *CCDC114* mutations appear to be fertile (Onoufriadis et al., 2013; Paff et al., 2013). It is suggested that its docking function may be replaced in sperm by the paralogue, *CCDC63* (Onoufriadis et al., 2013), which is orthologous to *wampa.* It seems likely that *CG14905/CCDC114* and *wampa/CCDC63* form functionally equivalent gene pairs in cilia and sperm respectively.

Why are there different ODA-DCs in *Drosophila,* and perhaps in other metazoans? One possibility relates to the fact that motile cilia and sperm flagella contain distinct ODA complexes, with different HCs and ICs, as has been shown for *Drosophila* (zur Lage et al., 2019) and humans (Thomas et al., 2020) (Fig. 8B). Thus, each ODA-DC may be adapted for recognising and binding different ODA subtypes. Consistent with this, human CCDC114 protein directly binds non-sperm HC, DNAH9. Alternatively, or in addition, ODA-DC differences may relate to differing modes of ciliogenesis. Chordotonal ciliogenesis is compartmentalised and requires IFT, while sperm flagellogenesis occurs within the cytoplasm independently of IFT (Han et al., 2003; Sarpal et al., 2003). Indeed, human TTC25 interacts with the IFT protein machinery (Wallmeier et al., 2016). Phylogenetic analysis of *CCDC151* homologues revealed a link between *CCDC151* genes and IFT (Jerber et al., 2014). *CCDC151* homologues are absent from organisms in which motile cilia assemble in an IFT-independent process.

## Acknowledgements

We thank Martyna Panasiuk for her help in the early stages of this project. We thank Bénédicte Durand and Jordan Raff for reagents. Stocks obtained from the Bloomington *Drosophila* Stock Center (NIH P40OD018537) were used in this study. This work was supported by a grant from the Biotechnology and Biosciences Research Council (BBSRC, BB/S000801) to AJ.

---

**Fig. S1.**
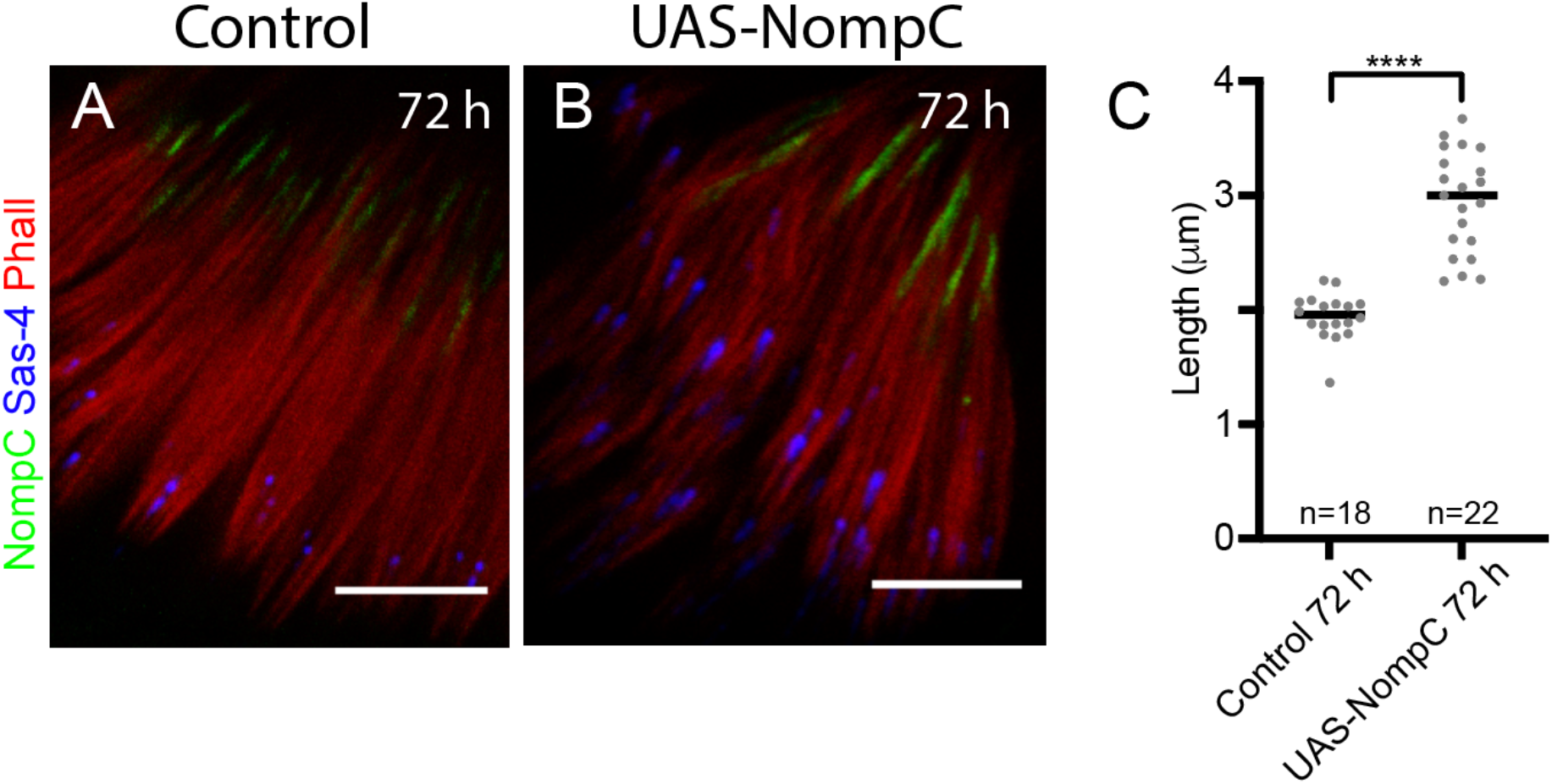
NompC overexpression does not strongly alter its localisation to the distal zone. (A,B) 72-h pupal antenna showing NompC (green), Sas-4 (blue), phalloidin (red). (A) Control *(scaGal4/+).* (B) NompC overexpression *(scaGal4, UAS-NompC),* showing longer extent of NompC labelling. (C) Graph comparing length of NompC labelling. Significance determined by two-tailed Mann-Whitney U test (****: P<0.0001). All scale bars are 5 μm.

**Fig. S2.**
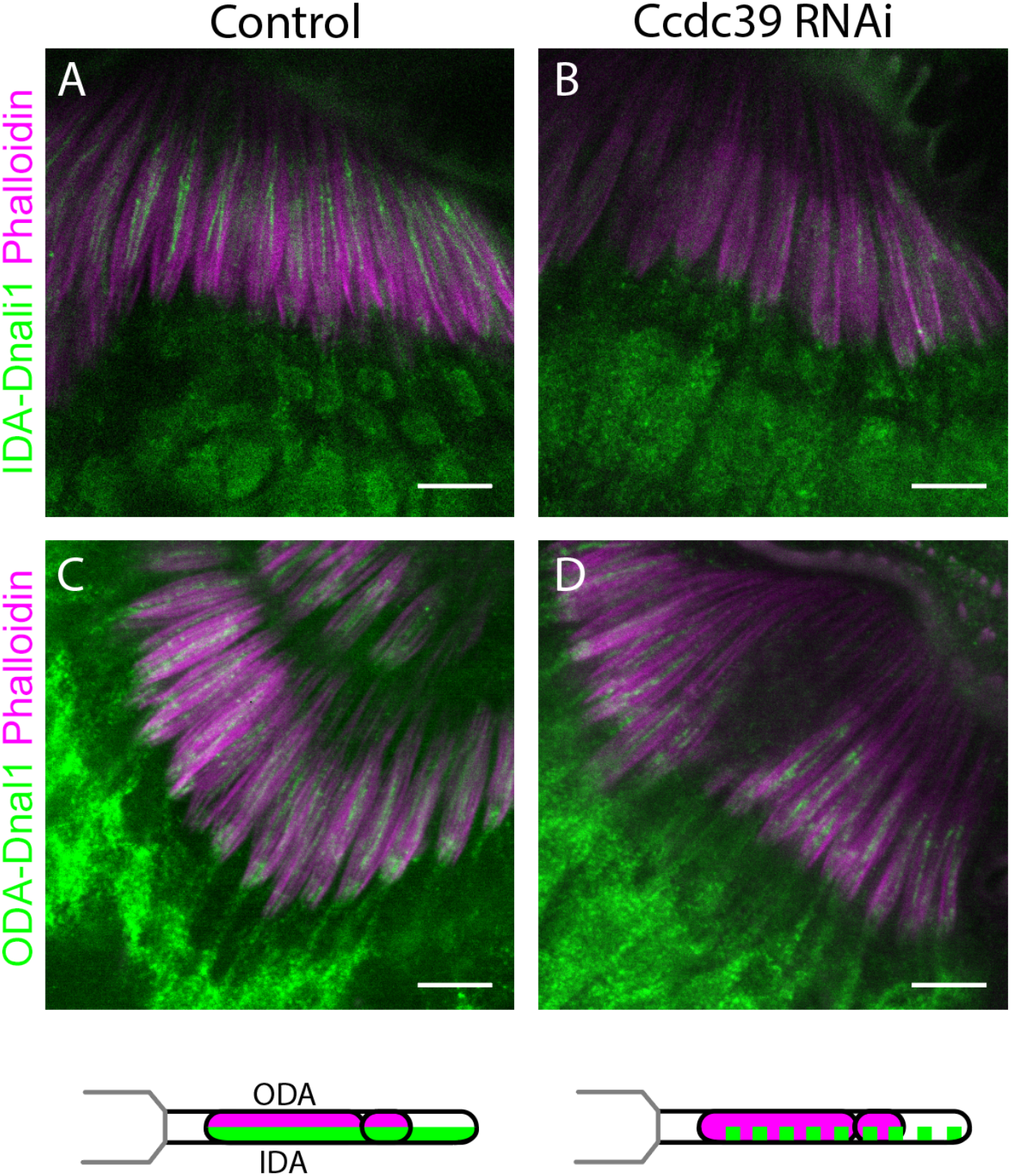
Knockdown of *Ccdc39* reduces localisation in the cilium IDA-Dnali1 but not ODA-Dnal1. 72-h pupal antennae. (A,B) Immunofluoresence for IDA-Dnali1-mVenus (green), phalloidin (magenta). (A) Control *(scaGal4/+).* (B) Ccdc39 knockdown *(scaGal4,* UAS-*Ccdc39*-RNAi). (C,D) Immunofluoresence for ODA-Dnal1-mVenus (green), phalloidin (magenta). (C) Control *(scaGal4/+).* (D) Ccdc39 knockdown *(scaGal4,* UAS-*Ccdc39*-RNAi). Below the panels is a schematic summary. All scale bars are 5 μm.

**Fig. S3.**
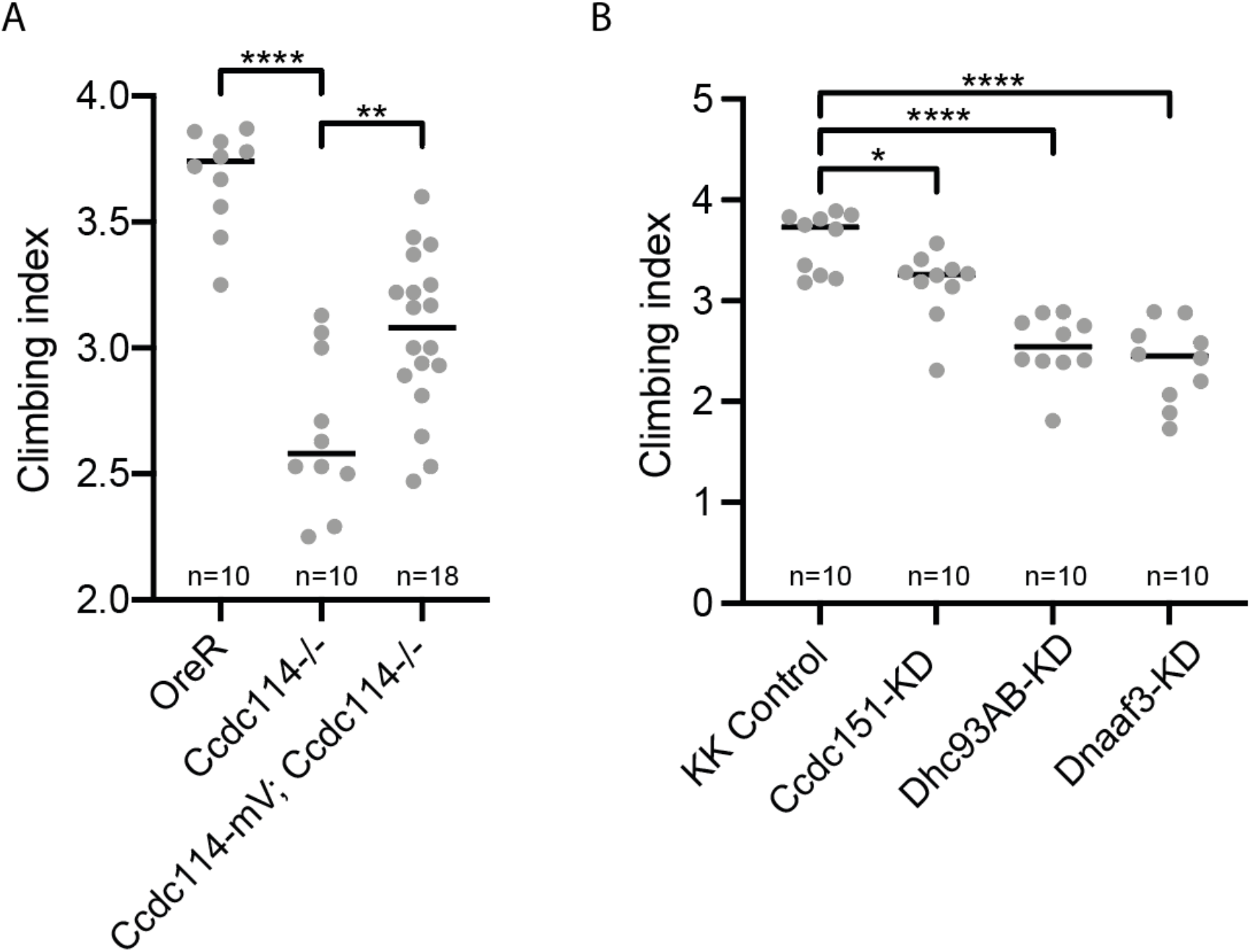
Climbing assays to test proprioceptive behaviour of flies. (A) *Ccdc114* homozygote flies perform significantly worse than wild-type flies (OreR) in a climbing assay (****: P<0.0001). Rescue of homozygotes by one copy of the Ccdc114-mVenus fusion gene improved their performance significantly compared to homozygotes (**: P=0.0027). Significance was determined by one-way Anova followed by Dunnett’s test for multiple comparisons. n = 10 batches of 15 flies (or 18 batches for rescue cross). (B) Knockdown of *Ccdc151* and *Dhc93AB (scaGal4,* UAS-RNAi) result in impaired performance in climbing assay compared to control *(scaGal4,* KK control line). For comparison, knockdown of *Dnaaf3* is also included, in which dynein arms are completely absent from the cilia (zur Lage et al., 2021). Significance was determined by one-way Anova followed by Dunnett’s test for multiple comparisons: *CG14127:* p = 0.0246; *Dhc93AB:* P < 0.0001; *Dnaaf3:* P<0.0001. n = 10 batches of 15 flies.

**Table S1.**
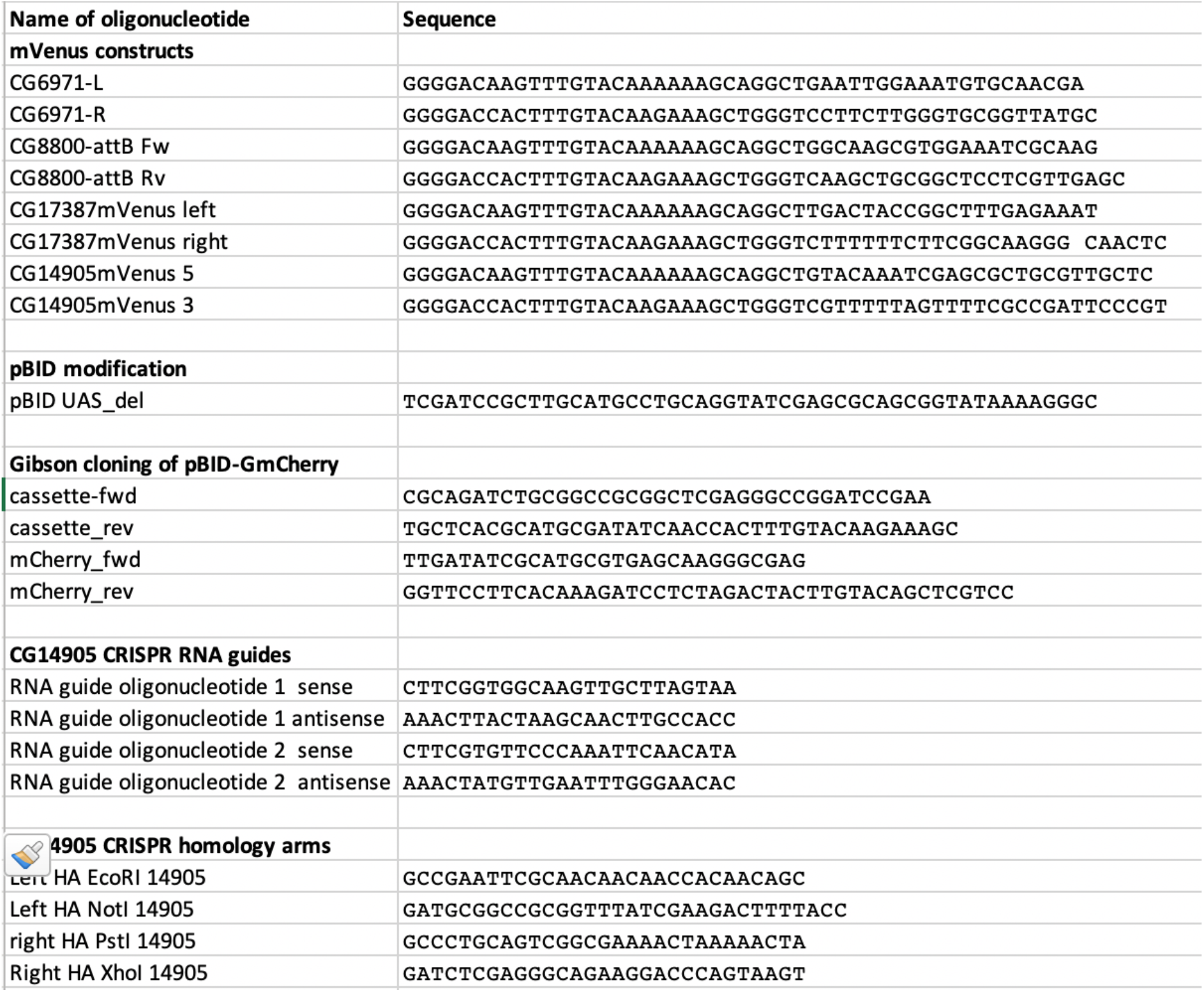
Oligonucleotides used in this study.

